# Deer slow down litter decomposition by reducing litter quality in a temperate forest

**DOI:** 10.1101/690032

**Authors:** Chollet Simon, Maillard Morgane, Schörghuber Juliane, Grayston Sue, Martin Jean-Louis

## Abstract

In temperate forest ecosystems, the role of deer in litter decomposition, a key nutrient cycling process, remains debated. Deer may modify the decomposition process by affecting plant cover and thus modifying litter abundance. They can also alter litter quality through differential browsing and affect decomposer ability by changing soil abiotic properties and the nature of decomposer communities. We used two litterbag experiments in a quasi-experimental situation resulting from the introduction of Sitka black-tailed deer *Odocoileus odocoileus sitkensis* on forested islands of Haida Gwaii (Canada). We investigated the effects of deer on decomposition through their impacts on litter quality and on decomposer ability. After one year, the effect of deer on litter quality resulted in a lower rate of mass loss in litter from litterbags. This mass loss mainly reflected a 21 and 38 % lower rate of carbon (C) and nitrogen (N) loss, respectively. Presence of deer resulted in lower decomposer ability for the rate of carbon loss, but not for nitrogen loss. The level of C loss after one year was 5% higher for litter decomposing on an island without deer. But the change in the rate of carbon loss explained by the effect of deer on decomposer ability was outweighed by the effect deer had on litter quality. Additional effects of deer on the decomposition process through feces deposition were significant but minor. These results question the role the large increase in deer populations observed in temperate forests at continental scales may play in broad scale patterns of C and N cycling.

## Introduction

### Deer and the functioning of temperate forests

Until recently, the role of large herbivores in nutrient cycling processes has been relatively neglected (Tanentzap & Coomes, 2012). In temperate forests from Europe and Eastern North America, dominated by coniferous or by broadleaved trees, this may be partly because ungulates, essentially deer, became largely missing as a result of hunting and/or loss of favorable land cover (McShea, Underwood, & Rappole, 1997; Apollonio, Andersen, & Putman, 2010). The extirpation of their natural predators, followed in the second part of the 20^th^ century, on both continents, by changes in hunting regulations and in land-uses, such as increased planting of winter crops by agriculture or, in some areas, farm abandonment and reversion to forests (see e.g. Côté, Rooney, Tremblay, Dussault, & Waller, 2004; Fuller & Gill, 2001; Milner et al., 2006) resulted in a dramatic rebound in deer populations that brought them back to the forefront of ecological thinking (Terborgh & Estes, 2013). The initial emphasis of research was on the consequences of deer recovery on forest vegetation, beginning with impacts on tree regeneration and growth (Gill, 1992), and, more recently, on aboveground understory community functioning (Horsley, Stout, & DeCalesta, 2003; Royo, Collins, Adams, Kirschbaum, & Carson, 2010), including cascading effects on different segments of the trophic network [invertebrates, birds (e.g. Chollet & Martin, 2013; Foster, Barton, & Lindenmayer, 2014)].

While our grasp of deer effects on forest aboveground communities has dramatically improved, their repercussions on belowground patterns and processes are still insufficiently understood (Bardgett & Wardle, 2003; Hobbie & Villéger, 2015). These belowground effects will be partly mediated by the effects deer have on litter decomposition and its pivotal role at the interface between aboveground primary production and belowground processes (Chapin, Matson, & Vitousek, 2011). In temperate forest ecosystems, contrary to grasslands or boreal forest, there are still only a few studies on how large herbivores affect litter decomposition.

### Deer and litter decomposition

Deer may modify belowground processes by affecting two of the main parameters that control decomposition: litter quality and decomposer ability (Prescott, 2010). Through plant removal, combined with selective foraging, deer modify plant community composition, plant stoichiometry, as well as the relative contribution of canopy and understory vegetation to litter composition (Côté et al., 2004). These changes in litter quantity and quality will affect decomposition processes and nutrient cycling (Bardgett & Wardle, 2003) .

Deer are also susceptible to modify belowground processes through the alteration of decomposer ability. This alteration can result from changes in edaphic properties such as increased soil temperature and salinity that follow exposure of bare soil after vegetation removal by browsing, or from soil compaction caused by trampling and its effects on soil water and oxygen content (Schrama et al., 2013). Deer also release dung and urine, a source of organic matter more easily decomposable than recalcitrant plant litter, and a source of inorganic nitrogen for soil decomposers that enhances their development (Sitters et al., 2017). These effects may affect the structure and functioning of decomposer communities [soil fauna (Andriuzzi & Wall, 2017) and microorganisms (Cline, Zak, Upchurch, Freedman, & Peschel, 2017; Eldridge et al., 2017)] with effects on the rate of litter decomposition (Handa et al., 2014).

Recent evidence indicate that decomposition occurs more rapidly when litter is placed under the plant species from which it originated (Gholz, Wedin, Smitherman, Harmon, & Parton, 2000; Ayres et al., 2009; Austin, Vivanco, González‐Arzac, & Pérez, 2014). This “home-field advantage” (HFA) is attributed to decomposer specialization. Home-field advantage may compensate the aforementioned potential changes in decomposition caused by deer. But studies explicitly testing this hypothesis are scarce, and provided contrasting results (see Olofsson & Oksanen, 2002; Penner & Frank, 2018).

There is recognition of the multiplicity of pathways through which deer may affect litter decomposition (see Bardgett & Wardle 2003 for a conceptual model), but our knowledge is mainly based on the independent study of each pathway, which led to apparent contradictions in results. To better identify the mechanisms behind the effect of deer on litter decomposition we designed a study that combined approaches able to disentangle the relative effects of these different pathways on the process.

### A quasi-experimental context

This study is part of a long term effort to use the introduction of Sitka black-tailed deer *Odocoileus odocoileus sitkensis* at the end of the 19^th^ century to the Haida Gwaii archipelago (British Columbia, Canada, see Golumbia, Bland, Morre, & Bartier, 2008) as an unplanned experiment on trophic interactions. Native to the coastal forests of British Columbia, Sitka black-tailed deer colonized most, but not all, islands, resulting in a quasi-experimental situation with, side by side, islands colonized by deer, and a limited number of small isolated islands never colonized. All these islands are forested. The occurrence of reference islands without deer made it possible to demonstrate that, on islands where deer were present, independent of island size, deer herbivory was the main factor structuring plant, invertebrate and songbird communities, overwhelming other biotic or abiotic factors (i.e. island area, soil and micro-habitat diversity (Table S1 in Supporting Information and Chollet, Baltzinger, Saout, & Martin, 2013; Gaston, Stockton, & Smith, 2006; Martin & Baltzinger, 2002; Martin, Stockton, Allombert, & Gaston, 2010). Recurrent experimental culls on some islands allowed to document the potential for recovery of the aboveground vegetation and songbirds (Chollet et al., 2016). As a result, islands can be segregated today into three categories of browsing histories and their associated vegetation patterns. Islands without deer, which are characterized by a diverse and lush understory vegetation dominated by broad leaved shrubs and ferns, producing a diverse and abundant litter. Islands where deer have been present for over 70 years, which are characterized by an open understory dominated by bryophytes where litter is dominated by conifer leaves (Martin et al., 2010; Stockton, Allombert, Gaston, & Martin, 2005). Finally, islands where deer have also been present for over 70 years but that have been subjected to recurrent deer culls over the past two decades. Their understory is characterized by an intermediate cover of vegetation (Chollet et al., 2016).

We used this extensive aboveground knowledge to design litterbag experiments based on these three browsing treatments, deer absent (no browsing, our reference), deer culled (intermediate browsing), and deer present (severe browsing), to analyze how the prolonged presence of abundant deer affected litter decomposition.

First, we assessed the effect of deer presence on litter decomposition at the scale of an island through a reciprocal litterbag translocation experiment involving litter representative of the three browsing treatments: “no”, intermediate” and “severe” browsing. The objective was to discriminate between the effects deer have on litter decomposition rate, either through their impact on litter quality or through their effect on decomposer ability (including effects on soil properties and on decomposer community composition), and this by explicitly including an assessment of home-field advantage by using the approach proposed by Keiser, Keiser, Strickland, & Bradford (2014). This approach allows to assess home-field advantage through the quantification of its role in decomposition rate relative to the role of litter quality per se (i.e. irrespective of the decomposition environment), and relative to the role of decomposer ability per se (i.e. irrespective of litter quality).

Second, we assessed how the deposition of high quality litter in the form of feces, affected the rate of litter decomposition at a narrow local scale by adding deer feces in a set of litterbags.

We predicted 1) that decomposition of litter collected from the deer-free island would be faster than decomposition of the litter collected from islands with deer (the effect of litter quality change); 2) that, in absence of home-field advantage, decomposition rates on the islands with deer would be higher than on the islands without deer due to an increase in decomposer ability in microbial communities in response to more recalcitrant litter (effect of change in decomposer ability); 3) that, presence of deer excrement would locally speed up the decomposition of plant litter; 4) that, the decomposition pattern observed on the islands where deer were culled should fall in between those observed on the islands with and without deer.

## Methods

### Study sites and plot selection

Haida Gwaii is characterized by a humid temperate-oceanic climate, with mean annual temperature of 8.5°C and precipitation that varies greatly from 1,350 mm on the east coast, where this study took place, to 7,000 mm on the west coast (Banner et al., 2014). The archipelago is covered by temperate rainforests dominated, at low elevation, by western hemlock (*Tsuga heterophylla*), western redcedar (*Thuja plicata*), and Sitka spruce (*Picea sitchensis*). The selected study sites belonged all to the Coastal Western Hemlock Wet Hypermaritime subzone [Biogeoclimatic Ecosystem Classification, code CWHwh1, (British Columbia Ministry of Forests, Lands, and Natural Resource Operations, 2010)] which covers 49% the archipelago, and ranges from sea-level to 350m in elevation (Banner et al., 2014). The bedrock geology of the selected study sites was volcanic and sedimentary, together with intrusions of granitic rock (Sutherland Brown, 1968). The soil type was organic and classified into the Folisol order (Soil Classification Working Group / Groupe de travail sur la classification des sols, 1998). We selected three islands in Laskeek Bay (52°53’12”N, 131°35’20”W, Fig. S1 in Supporting Information). We chose sampling and experimental sites with similar parent material, and that were representative of the patterns of deer impacts we documented at the scale of the archipelago (Chollet, Bergman, Gaston, & Martin, 2015; Martin et al., 2010) (see Table S1 in in Supporting Information for a synthesis of previous studies). The three islands were Low Island, 9.6 ha, that had never been colonized by deer, Louise Island, 25,000 ha, that has had deer for over 70 years (Vila, Torre, Guibal, & Martin, 2004) and had a current deer density estimated at 30 deer / km², and Reef Island, 249 ha, that had also been colonized by deer more than 70 years ago, but its deer population had been regularly culled between 1997 and 2010. At the time of study Reef Island had a deer population density estimated at about 15 deer / km² and a partially recovered understory vegetation (Chollet et al., 2016). Low, Reef and Louise Islands were therefore representative of three distinct deer herbivory treatments: absence of current or past browsing, intermediate browsing pressure, and severe browsing pressure, respectively. Our emphasis on selecting sites similar in parent material and similar in forest types prevented us to control also for island size. To take this into account we selected all sampling plots in the coastal area on the three islands. On each island we established fifteen 10 m × 10 m forest interior plots, leading to a total of 45 plots. Adjacent plots were separated by at least 100 m.

### Above and belowground characteristics in relation to deer presence

In each plot we sampled the vegetation by estimating the percent cover of vascular plants and bryophytes using the Londo scale (Londo, 1976). We measured soil bulk density at the surface of the soil with five replicate measures per plot. For this, we collected soil with a 5.4 cm depth × 4.1 cm diameter (71.29 cm3) copper core hammered into the soil using a mallet. We took care to not change the structure of the soil while sampling. We removed any coarse woody debris from core samples and subtracted their volume from the volume of the core. We then dried soil at 105°C for 24h to obtain a value for bulk density expressed as g of dried soil per cm^3^ of fresh soil. We used data on soil pH, C:N and organic horizon depth collected in the course of a sister study in plots located in the same area on these same islands (Maillard et al. unpublished data). This data was collected from five plots on Low Island, five plots on Louise Island and six plots on Reef Island. We sampled soil within these plots with a 2.5 cm diameter × 30 cm depth core collecting approximately 100 cores per plot. They were mixed and sieved with a 5 mm sieve to ensure homogenization as recommended for soil with high content of organic matter (Haynes & Swift, 1990). We measured soil pH in a 0.01 M CaCl2 solution using a 1:10 ratio (air dried soil: solution). We determined Soil C:N ratio from 3 mg of freeze-dried and ground soil using an Elementar Vario El Cube Analyzer. In each plot we measured the depth of the soil’s organic horizon from a soil pit dug within the plot.

### Experimental design and protocol

We measured litter decomposition rates using the litterbag method. We made 15 cm * 15 cm bags using polypropylene mesh with two different mesh sizes. We used the litterbags with a 0.2 mm mesh size to target the decomposition solely due to soil microfauna and microorganisms. We used litterbags with a 3.7 × 4.45 mm mesh size to assess the additional effect of mesofauna and macrofauna on litter decomposition. To obtain our litter samples, we collected summer senescent leaves from plant species with a percent cover on the plots greater than 5 %. In total, we sampled 18, 20 and 17 plants species respectively on the island with no browsing, on the island with an intermediate level of browsing, and on the island with a severe level of browsing (Table S2 in in Supporting Information). We dried these litter samples at 30°C for a week before using them.

We developed two complementary experiments in order to study the various mechanisms by which deer could affect the decomposition process.

#### Experiment 1

To investigate litter decomposition rate in relation to the three browsing treatments we produced, for each plot, three identical litterbags for each of the two mesh-sizes. Each of these six litterbags contained, in the same proportion as in the plot, plant material from all plant species covering more than 5 % of the plot area. We fixed the total mass of litter per litterbag at 4 g. Hence, the mass of litter from each plant species in a given litterbag was calculated according to its relative abundance in the plot. For each mesh size we placed one of the three litterbags on the plot the litter came from (“home”), and placed the two remaining bags on one plot on each of the two remaining islands (“away”). This translocation allowed us to independently test for the relative effects of home-field advantage, of decomposer ability (soil properties and decomposer community composition), and of litter quality on litter decomposition (Fig. S1).

#### Experiment 2

To investigate the influence of deer feces on litter decomposition rate within a litterbag we used a standardized litter. We chose litter from one of the dominant tree species on all islands, *P. sitchensis*. To avoid any biases from potential inter-treatment differences in spruce litter quality, we used, for this set of litterbags, a mix of *P. sitchensis* litter collected from all three islands. In order to standardize feces quality we collected fresh deer feces from another island with deer, also situated in Laskeek Bay (East Limestone, 48 ha). We used this material to place on each of the 45 plots one fine-mesh litterbag containing 2 g of deer feces and 2 g of the standardized litter and two fine-mesh litterbags containing controls, one filled only with 5 g of deer feces and one filled only with 2 g of *P. sitchensis* litter. We repeated this with coarse-mesh litterbags.

Thus, to implement these two experiments, we placed a total of 12 litterbags on each plot including 6 fine-mesh litterbags [3 for experiment 1 (1 home and 2 away) and 3 for experiment 2 (1 feces only + 1 *P. sitchensis* only + 1 with feces and *P. sitchensis*)] and a similar set of 6 litterbags for the coarse-mesh litterbags. As a result we had a total of 540 litterbags (12 bags * 45 plots) for the full design. All litter bags were identified and numbered with aluminum tags and placed randomly on the surface of the forest floor on each plot. We used U pins at each corner of the bag to hold them in place. We placed litterbags on the three islands in July 2017, and collected them one year later in July 2018. The remoteness of the islands and associated logistics prevented designing an experiment with a partial collection of litterbags during the year to better assess the kinetics of decomposition. After collection (all but 35 bags were retrieved) we dried the litterbag contents at 70°C for 48h prior to weighing the contents and then performing chemical analyses.

### Mass, Carbon and Nitrogen loss in litterbags

To assess litter mass loss over a year in each litterbags we subtracted the final mass of the bag’s content from its initial mass. To assess C and N loss over a year, we calculated C and N contents of samples with an Elementar Vario El Cube Analyzer (Elementar, Langenselbold, Germany) using 3.5 mg of ground material. We first calculated the initial C and N contents of dried litter using eight individuals of each plant species (vascular and bryophytes) from each island/browsing treatment. Based on these values, and on the relative proportion of each litter in the bags, we calculated the initial C and N content for each litterbag. We also measured initial C and N contents of eight deer pellet groups that were previously dried at 70°C for 48h.

At the end of the experiment we finely ground the dried litter from the fine-mesh litterbags only, to remain within our budget limitations, and measured C and N content. We calculated carbon and nitrogen loss by subtracting the amount of carbon and nitrogen remaining in litter after one year of decomposition from the initial estimates based on our calculations of initial C and N content of the litter material.

### Statistical analysis

In order to evaluate the effect of deer herbivory on plant community composition we used a Correspondence Analysis on our data of plant species cover per plot, and performed a between class analysis (Dray & Dufour, 2007). We evaluated the significance of the class effect with a permutation test. To assess the initial litter C:N ratios at each plot we calculated the Community Weighted Mean (CWM) of this initial litter C:N ratio using the formula

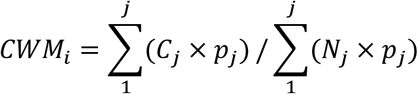

where *i* represents the plot, *j* the plant species on this plot, *C*_*j*_ and *N*_*j*_ the C and N content of the corresponding litter, and *p*_*j*_ the relative abundance of the corresponding plant species on the plot.

We used one way ANOVA with permutation tests to compare litter CWM C:N ratio, soil bulk density, soil pH, soil C:N ratio and organic horizon depth among the three islands. We used the multiple comparison post-hoc test with the function *kruskalmc* from the package pgirmess.

To assess differences in the rate of litter mass loss and of C and N loss in both experiments, we calculated the percent differences among treatments in litter mass loss and in C and N loss relative to the no browsing treatment using the formula:

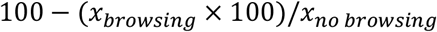

Where *x*_*browsing*_ is the litter, C, or N mass loss in litterbags from either the intermediate or the severe browsing treatment, and *x*_*no browsing*_ the litter, C, or N mass loss in litterbags from the no browsing treatment.

#### Analyses of decomposition experiment 1

We used a two way ANOVA to compare litter mass loss, carbon loss and nitrogen loss from litter after one year among the three origins of litter (from island with no, intermediate or severe deer browsing) and among treatment categories (island with no, intermediate or severe deer browsing).

To disentangle the relative importance of the two main ways deer may modify C and N decomposition, namely litter quality and decomposer ability, we used the Decomposer Ability Regression Test proposed by Keiser et al. (2014) using SAS 9.4 (SAS Institute, Cary, NC). This method statistically discriminates among effects of litter quality (here defined as how rapidly a litter is decomposed regardless of decomposition site), ability [i.e. how rapidly a litter is decomposed at one site regardless of litter quality (includes the effect of soil abiotic conditions and the ability of the decomposer communities)] and home-field advantage [i.e. the acceleration of litter decomposition when litter is placed in the site it comes from (home) and where it can potentially benefit from a local specialization of the decomposer community]. The regression model defines the rate of decomposition of observation i (Y_i_) by three parameters: litter quality (Litter_l_), soil ability (Soil_s_), and HFA (Home_h_) which are dummy variables that equal 1 or 0, respectively, depending on the presence or absence of the litter mixture, soil community and home combination (in observation i). The parameters to be estimated are β_l_, γ_s_ and η_h_ (Keiser et al., 2014). The average decomposition across all data (i) in a dataset, after controlling for litter, soil and home combinations, is represented by the intercept (α), and the error term is defined by ε. The β_l_ and γ_s_ are restricted to prevent collinearity.

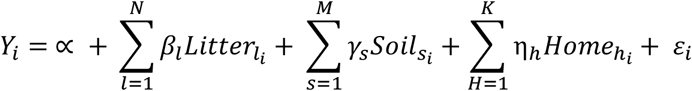

Compared to the classically used Structural Equation Modeling (SEM), the Decomposer Ability Regression Test offers the additional possibility to explicitly test home-field advantage and maximizes the information extracted from the litter transplant experiment (Keiser et al., 2014).

To explore the reasons for differences in litter quality caused by deer we performed linear models between C or N loss and litter CWM C:N ratio.

#### Analyses of decomposition experiment 2 (feces manipulation)

We used a two way ANOVA to compare mass loss, C loss, and N loss of *Picea sitchensis*, feces and the combination of both.

For all analyses in which homoscedasticity and normality of the distribution of the residues were not respected, we used ANOVA with permutation tests instead of classical ANOVA (lmPerm package, (Wheeler, 2010).

We used the R 3.4.1 environment (R Core Team, 2017) for all statistical analyses (except Decomposer Ability Regression Test).

## Results

### Deer modify aboveground and belowground characteristics

The first axis of the Correspondence Analysis significantly discriminated the plant species composition and abundance in the plots according to the intensity of deer browsing (Fig. 1A, Monte-Carlo permutation test: p<0.001). In the absence of deer, vegetation cover was higher and there was greater shrub diversity (Table S2 in Supporting Information). The vegetation from plots with severe deer browsing was characterized by a high cover and diversity of bryophytes (Fig. 1A). Plots under intermediate deer browsing showed intermediate plant species diversity and cover. We found no significant difference in the initial C:N ratio of the plant litter among deer browsing treatments (Fig. 1B, p-value = 0.2). Soil bulk density was significantly higher on plots from islands with deer (Fig. 1C, p-value < 0.001). Soil pH decreased significantly with increasing deer browsing pressure (Fig. 1D and F, p-value = 0.037 and 0.005 respectively, however statistical power was not sufficient to discriminate which treatment is different in the post-hoc test). Soil C:N was not significantly different among treatments (Fig. 1E, p-value = 0.32). Depth of the organic horizon measured in the plots decreased by 44 % with increasing browsing intensity.

**Figure 1.**
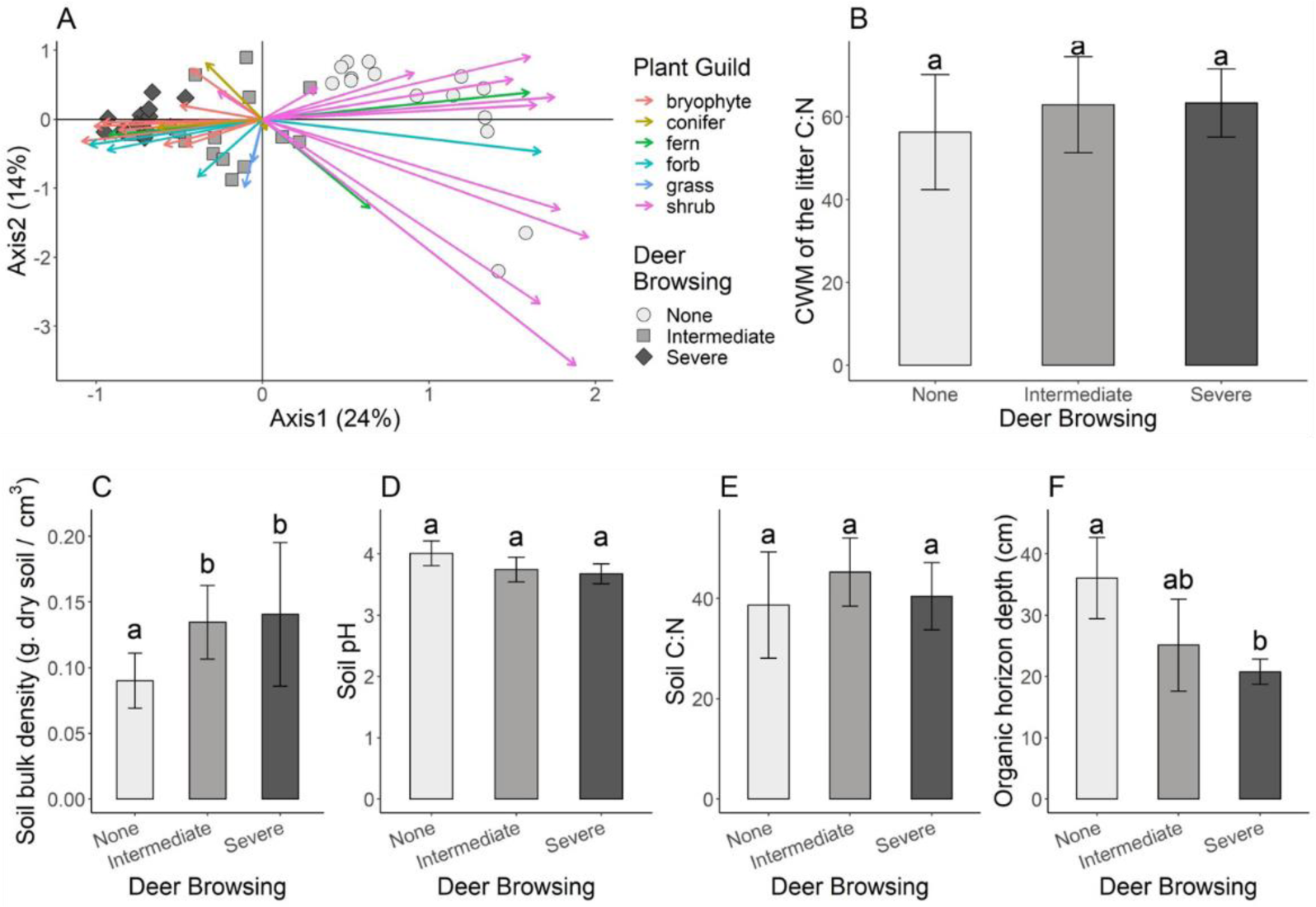
Effect of deer herbivory on aboveground (A, B) and belowground (C to F) parameters. Shades of dots and barplots represent the deer browsing treatment with: light grey = no browsing (deer absent), grey = intermediate (deer present for over 70 years but exposed to significant culls between 1997 and 2010) and dark grey = severe browsing (deer present for over 70 years and not exposed to hunting). Small letters on each barplot indicate differences tested by non-parametric post-hoc test. Panel A - Correspondence Analysis on the vegetation data collected at each plot. Dots, squares and lozenges represent the coordinates of the plots from the islands with no browsing, intermediate browsing and severe browsing, respectively. Arrows indicate the species contributions to axes (one arrow per species). Plant species are classified according to their functional group; Panel B – Community Weighted Mean (CWM) of C:N ratio of the plant community; Panel C-Soil bulk density; Panel D -Soil pH; Panel E-Soil C:N ratio; Panel F - Organic horizon depth.

### Litter mass loss in experiment 1

Litter mass loss was highest in litterbags with litter from the island with no deer and lowest in litterbags with litter from the island with severe browsing pressure. In fine-mesh litterbags average litter mass loss was 55% in litter from the islands without deer, 43% for litter from islands with intermediate browsing pressure and 34% in litter from islands with severe browsing pressure (Fig. S2A in Supporting Information). This pattern hold whatever the island category litterbags were placed on (Fig. S2A). Variation due to the context in which a given category of litter was placed had only little influence, there was no significant effect of home-field advantage (Figs. S2A and S2B). Decomposers were not more efficient in decomposing litter when it originated from their own environment rather than from other sites. Mass loss was significantly affected by the place of decomposition (Table S3, F = 113.36, p-value = 0.05). This pattern was due to a significantly better ability of the micro-fauna and microorganisms from plots on the island without deer to decompose litter (Figs S2A and S2C, p-value = 0.0045). Litter quality, understood here as the rate of decomposition independent of decomposer ability and home-field advantage [calculated using the method developed by Keiser et al., (2014)], was the main driver of carbon loss (Fig.S2D). Litter mixes originating from the island with no deer had the best quality index (highest loss after one year, first three bars on Fig. S2A), followed by the litter mixes originating from the island with intermediate deer browsing and then by litter mixes originating from the island with severe deer browsing (Figs. S2A and S2D).

In coarse-mesh litterbags the overall patterns of decomposition was similar to those from fine-mesh litterbags (compare Figs. S2A to S2D and Figs. S2E to S2H in Supporting Information), although variability among plots within sites was greater (Figs. S2A and S2E). As a result, mass loss in litter mixes originating from islands with intermediate and severe deer browsing were respectively 25% and 39% lower than in litter mixes originating from the plots on the island without deer (p-value < 0.001) (Fig. S2C). There was no evidence for home-field advantage in large mesh litter bags (Fig. S2F). Place of decomposition significantly affected litter mass loss (Figure S3, F = 4.54, p-value = 0.01). The ability of the micro-fauna and microorganisms to decompose litter was significantly lower for litter placed in plots exposed to severe browsing (Fig. S2G, p-value = 0.009). As observed for fine-mesh litterbags, litter quality had a significant effect on litter decomposition in coarse-mesh bags (Fig. S2H, p-value < 0.001).

### Carbon and Nitrogen loss in fine-mesh litterbags in experiment 1

The loss of carbon in the litterbags after one year was highest for litter representative of the vegetation on islands without deer and lowest for litter representative of the vegetation on islands with the most severe browsing pressure (Fig. 2A, F = 108.78, p-value = < 0.001) a pattern consistent with the pattern of litter mass loss. When compared to the carbon loss observed in litter originating from the island with no deer, carbon loss after one year was 12% lower in litter collected from the island with intermediate browsing, and 30% lower in litter collected from the island with severe browsing, this independently of the incubating (i.e. island) context (Fig. 2A). Home-field advantage was not significant (Fig. 2B), indicating that decomposers were not more efficient in decomposing litter carbon when it originated from their own environment rather than from other sites. The ability of the micro-fauna and microorganisms to decompose carbon was significantly higher for litter placed in plots without deer than in plots on islands with deer (Fig. 2C, p-value = 0.008). Indeed, carbon loss after one year was 5% lower in litterbags incubated in plots on the island with severe deer browsing than carbon loss observed in plots from the island without deer. Litter quality, understood here as the rate of decomposition independent of decomposer ability and home-field advantage [calculated using the method developed by Keiser et al., (2014)], was the main driver of carbon loss. Litter mixes originating from the island with no deer had the best quality index (highest loss after one year), followed by the litter mixes originating from the island with intermediate deer browsing and then by litter mixes originating from the island with severe deer browsing (Figs 2A and 2D).

**Figure 2.**
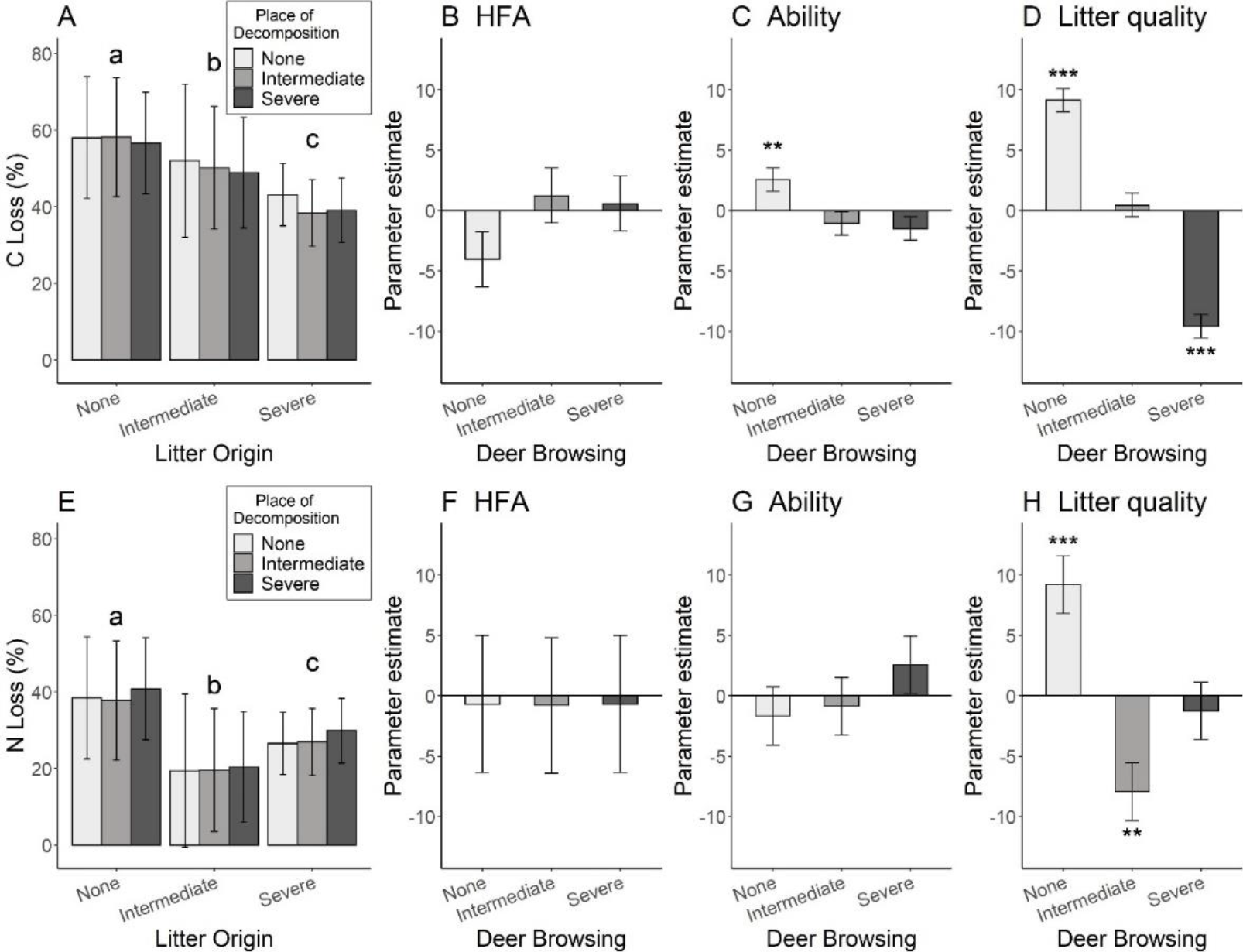
Decomposition rate of the plant community litter among deer browsing categories for carbon (top) and nitrogen (bottom) in fine-mesh litterbags in the translocation experiment. Shades of barplots represent the deer browsing intensity with: light grey = no browsing (deer absent), grey = intermediate (deer present for over 70 years but deer density reduced by culls between 1997 and 2010) and dark grey = severe browsing (deer present for over 70 years but not exposed to hunting, highest deer density). Asterisks indicate estimates significantly different from zero with *<0.05, ** <0.01, ***<0.001. Fine letters in each bar plots indicate differences tested by post-hoc test. Panel A and Panel E represent carbon and nitrogen loss after one year among treatments respectively with bars grouped according to litter origin and shades corresponding to the category of deer browsing of the location where the litter bags were placed. Panel B-D and F-H represent the parameter estimates (± SE) calculated using the Decomposer Ability Regression Test proposed by Keiser et al. (2014).

Litter mixes from the island without deer had significantly higher nitrogen loss than litter mixes from islands with deer (Fig. 2E). However, unlike C loss, nitrogen loss was lower for litter mixes originating from the island with intermediate deer browsing than that originating from the island with severe deer browsing (45% and 30% respectively, Fig. 2E, F = 17.53, p-value < 0.001). We detected no home-field advantage for N loss (Fig. 2E). In addition, none of the decomposer communities were better at decomposing and releasing nitrogen (Fig. 2F). Similarly than for carbon, litter quality (sensu Keiser et al. 2014) was the main driver of nitrogen loss in litter bags. However, conversely to carbon, the lowest rate of N loss after one year was observed in litter from the island with intermediate browsing pressure (Figs 2E and 2H).

Carbon and nitrogen loss in litterbags after one year were significantly and negatively related to the initial C:N ratio Community Weighted Mean (CWM) of the litter (Fig. 3). For litter carbon loss, the initial litter C:N CWM explained only 10% of its variation (Table S2). Conversely, litter nitrogen loss was strongly linked to the initial litter C:N CWM, which explained 50% of its variability (Table S4 in Supporting Information).

**Figure 3.**
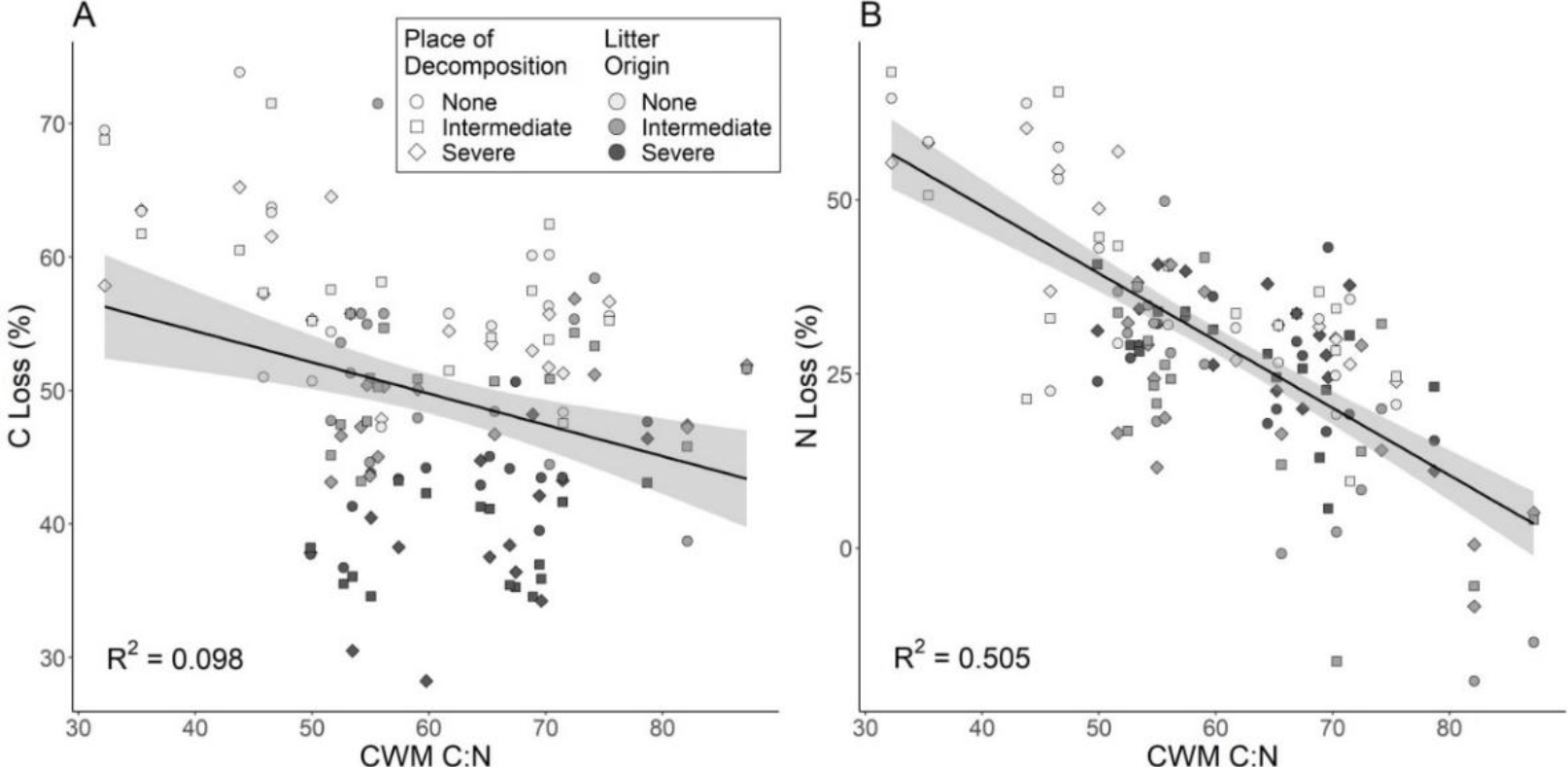
Linear regression of carbon (left) and nitrogen (right) loss variation with plant C:N Community Weighted Mean. Shades of dots represent the deer browsing intensity on the island where the litter came from, with: light grey = no browsing (deer absent), grey = intermediate (deer present for over 70 years but culled, lower deer density) and dark grey = severe browsing (deer present for over 70 years but not exposed to hunting, highest deer density). The shape of the symbols refer to the browsing category of the island where we placed the litterbags. Details on regressions models are given in Table S2.

### Feces decomposition in experiment 2

In fine-mesh litterbags place of decomposition had no effect on feces mass loss (Fig. S3A). In coarse-mesh litterbags mass loss from feces after one year was 15% higher on islands with deer than on islands without deer (Fig. S3B, p-value < 0.001). The addition of feces enhanced the mass loss in *P. sitchensis* litter by 29% in fine-mesh (p-value = <0.001, Fig. S3C) and by 20% in coarse-mesh litterbags (p-value = 0.047, Fig. S3D), on island with intermediate or no browsing (Fig. S3D). There were no differences among treatments in carbon and nitrogen loss from feces after one year (F = 1.386, p-value = 0.26 and F = 0.416, p-value = 0.66 respectively, Figs. 4A and B). Feces addition significantly increased the C and N loss in *P. sitchensis* litter after one year, by 31% for carbon and 47% for nitrogen (F = 175.62, p-value < 0.001 and F = 66.39, p-value < 0.001 respectively, Fig. 4C and D). The ability of the decomposer community (i.e. decomposition place) had no effect on carbon loss (F = 0.752 and p-value = 0.47, Fig. 4C). However, for nitrogen loss in *P. sitchensis* litter after one year, the presence of deer feces significantly improved the decomposition ability of the decomposer community in the plots from the islands with deer (F = 20.10, p-value < 0.001, Fig. 4D).

**Figure 4.**
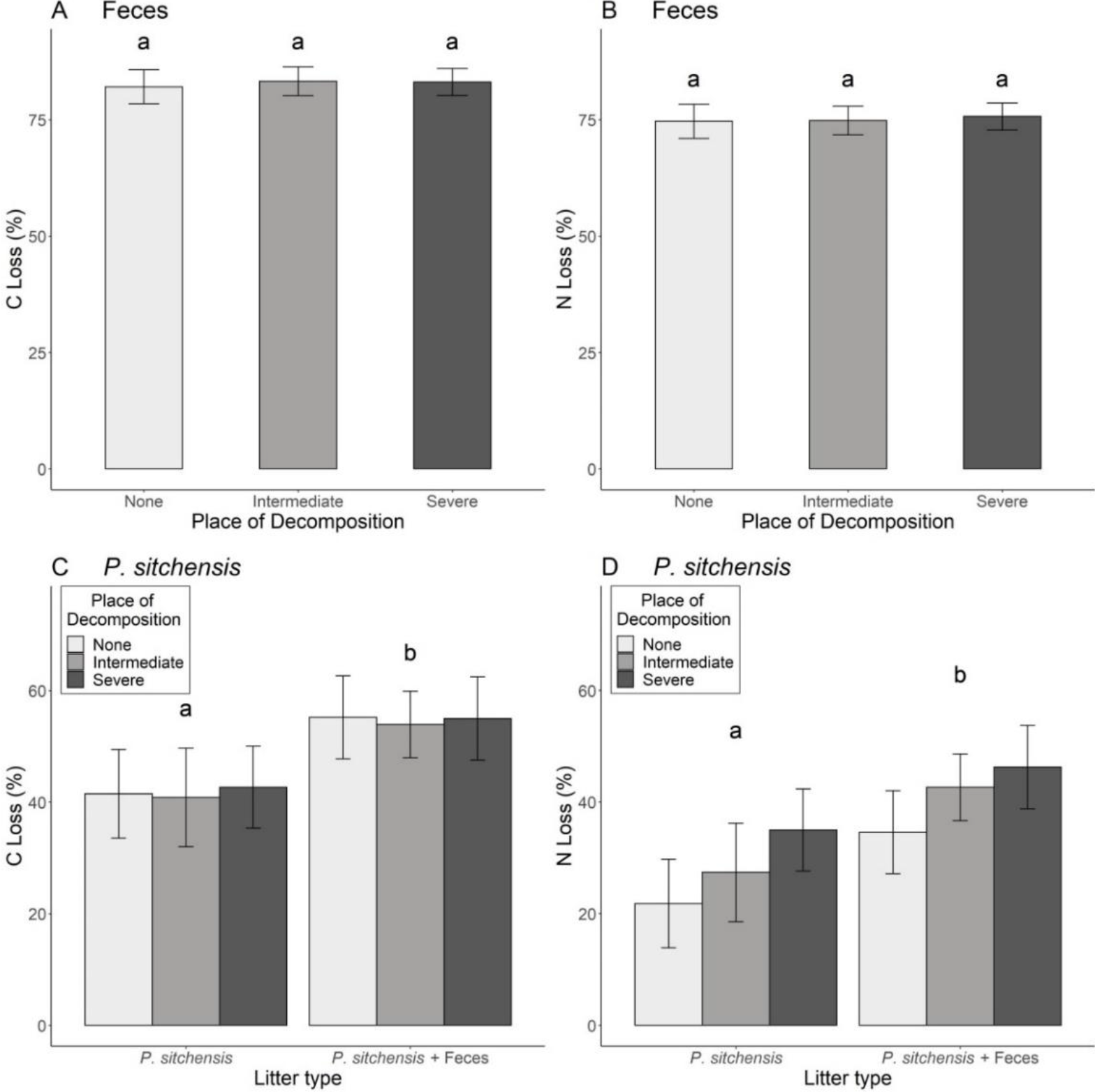
Carbon and nitrogen loss in feces in relation to browsing treatment (top) and effect of feces addition on carbon and nitrogen loss in *Picea sitchensis* litter (bottom) in fine-mesh litterbags. Shades of barplots refer to the deer browsing category of the place of decomposition with: light grey = no browsing (deer absent), grey = intermediate (deer present for over 70 years but not exposed to hunting, highest deer density) and dark grey = severe deer browsing (deer present for over 70 years but not exposed to hunting, highest deer density). Panel A – Carbon loss in feces; Panel B – Nitrogen loss in feces; Panel C – Carbon loss in *P. sitchensis* litter with and without the addition of feces; Panel D – Nitrogen loss in *P. sitchensis* litter with and without the addition of feces. Small letters above each barplots indicate differences tested by post-hoc test.

## Discussion

### Deer slow down decomposition through modification of the understory plant communities

To our knowledge, our study is among the few studies on the effect of large forest herbivores on decomposition processes that have attempted to dissociate the relative effect of changes in plant community composition (community litter quality) from the effects of changes in abiotic soil properties and decomposer community (decomposer ability, Andriuzzi & Wall, 2017). Previous research focused on decomposer ability and, when integrating litter quality, considered only the plant specific responses to herbivory (i.e. changes in plant chemical composition associated to browsing), but neglected the change in plant species composition and relative abundance. In our study, we demonstrated that herbivores change the overall quality of litter reaching the forest floor, and that this is the overriding factor governing litter decomposition, rather than soil properties, composition of the soil decomposer community, or home-field advantage (Fig. 2). This difference in overall litter quality was not exclusively attributed to a modification of the litter C:N CWM (Figs. 1B & 3). This suggests that other parameters of litter quality not measured in our study, such as lignin or anti-herbivore compounds, were also implied in overall litter quality decline in presence of deer. Thereafter we refer to the term ‘litter quality’ as a composite variable calculated as proposed by Keiser et al. (2014), and representing the rate of decomposition independent of decomposer ability and home-field advantage. The major shift in litter quality caused by deer browsing resulted, after one year, in an overall reduction in litter mass loss in presence of deer. This translated after one year into a 25% lower carbon and a 27% lower nitrogen loss in litter from the island with severe browsing compared to C and N losses in litter from the island with no browsing (Figs. 2A and 2E). This strong control of litter quality on C, and especially on N loss, contrasts with the previous assumption that vegetation changes may affect nitrogen and phosphorus dynamics less than the dynamics of carbon (Bryant, Chapin, & Klein, 1983; Wardle, Bonner, & Barker, 2002).

The prevailing importance of change in litter quality that resulted from deer herbivory on decomposition in the temperate forests we studied is in agreement with the microcosm study of Harrison & Bardgett (2003) who showed that decomposition of birch (*Betula pubescens*) litter, originating from inside deer exclosures (unbrowsed) in the Scottish Highlands decomposed faster than litter from outside of the exclosures (browsed), irrespective of the origin of the soil (inside or outside of exclosures). Conversely, Olofsson & Oksanen (2002), in a field translocation experiment assessing the decomposition of four plant species dominating the vegetation of lightly and heavily grazed tundra demonstrated a positive effect of reindeer herbivory on decomposition rate.

The dramatic change in litter quality found in our study is the result of an alteration in the understory plant community (Figs. 1A, 2D and 2H). Intense and prolonged deer browsing dramatically changed the understory plant composition and cover, resulting in an up to 90% reduction in understory shrub cover (Table S1 Fig. 1A), and in a shift in litter quality. We interpret this change in litter quality on islands with as the main cause for the dramatic reduction in litter decomposed after one year (Figs S2D and S2H and Figs. 2D and 2H). These modifications not only confirm the severe impact of deer on the understory vegetation of Haida Gwaii (Chollet, Baltzinger, Ostermann, Saint-André, & Martin, 2013; Martin et al., 2010) but are consistent with results in other temperate forests (Côté et al., 2004; Boulanger et al., 2018).

We found that the reduction in litter quality, and the associated modifications in the decomposition pattern, were partly driven by the variation in the litter CWM C:N ratio (Fig. 3) although we found no overall difference in litter CWM C:N ratio among islands (Fig. 1B). The decline in litter quality affected carbon and nitrogen cycles differently. For carbon, litter quality decreased as deer browsing intensity increased (Fig. 2D). For nitrogen, litter quality was poorer in the intermediate deer browsing than on the island with severe browsing (Fig. 2H). The decline in litter C loss could be explained by the shift from an understory dominated by more decomposable species (shrubs) towards an understory of less decomposable species (conifers and bryophytes) as the level of deer browsing intensity increased. Conifers and bryophytes are known to have slow decomposition rates due to low N content and high concentrations of structural carbohydrates and aromatic compounds (Cornwell et al., 2008; Turetsky, Crow, Evans, Vitt, & Wieder, 2008). The presence of these secondary compounds may largely explain the lower decomposition of the litter from islands with deer and thus the slight effect of CWM of litter C:N (≈10% of variation explained, Fig. 3). The contrasting result we obtained for N loss from litter after one year suggests that the vegetation shift caused by deer had different consequences for nitrogen mineralization. Although there was no overall significant difference in CWM litter C:N ratio among deer herbivory treatments, the intermediate treatment had the highest values of C:N (Fig. 3), which explains the lowest N loss values observed in this treatment (≈50% of the variation is explained by CWM litter C:N).

### Deer also modify decomposer ability

Although the change in litter quality (sensu Keiser et al. 2014) caused by deer herbivory was identified as the main driver of the decomposition process, several other changes in the soil decomposer communities affected nutrient cycling. Decomposers from the island without deer had a greater ability to decompose the carbon present in litter, but not nitrogen (Figs. 2C and 2G). The contrast between C and N decomposition among islands when using fine–mesh litterbags, and the similarity in C and N decomposition when using coarse -mesh litterbags (Fig. S2), suggests that the observed decomposition patterns are more likely explained by biotic differences in soils (i.e. differences in decomposer community) than by the effect of abiotic modifications such as higher soil compaction (Fig. 1C). A possible explanation for the observed contrasts in litter decomposition may be a switch in the bacterial:fungal ratio in presence of deer. In fact the disappearance of base-rich shrubs and their replacement by species with high concentrations of phenolic compounds (e.g. bryophytes) as a result of deer browsing may have increased the dominance of fungi which require less calcium and magnesium for growth (Prescott, 2010). This change in decomposer community structure would favor the formation of a mor humus, in which up to 30% of the litter mass is converted to humus rather than decomposing (Prescott, 2010). In addition the dramatic reduction in shrub cover may have reduced the root exudate which stimulate bacterial activity (Ekberg, Buchmann, & Gleixner, 2007). Contrary to carbon, the ability of decomposers to decompose nitrogen in litter did not vary among islands with different patterns of deer herbivory (Fig. 2G). This may be explained by the selection of microorganisms better able to exploit N in environments where this element is the most limiting (“nitrogen mining hypothesis”, Craine, Morrow, & Fierer, 2007) compensating for the switch in bacterial:fungal ratio. This hypothesis is supported by our control experiment, where we used a standardized quality of litter *(Picea sitchensis*), and found a greater ability of decomposers to decompose N in litter samples incubated on sites with deer (Fig. 4D).

Interestingly, we also found that the inclusion of the soil macro and mesofauna (hereafter fauna) in litter decomposition via the use of coarse-mesh litterbags affected litter mass loss (Fig. S2E). Specifically, we found that litter decomposition was reduced on the island with the highest deer density, suggesting a negative effect of high deer density on the faunal decomposer communities. Previous studies documented negative effects of large herbivores on the abundance and diversity of the soil fauna (see review by Andriuzzi & Wall (2017)), but the consequences on litter decomposition were not studied. This negative effect could be due to a reduction in both the abundance and the activity of the soil fauna through several mechanisms. Directly, through soil trampling by deer which might reduce soil fauna habitat through soil physical compaction and its reduction of soil pore size (Beylich, Oberholzer, Schrader, Höper, & Wilke, 2010). In addition, the reduction of litter quality by deer might be responsible for an indirect slowing down of soil faunal abundance and activity for which litter quality is known to be a controlling factor (García‐Palacios, Maestre, Kattge, & Wall, 2013; Hendriksen, 1990). As most previous studies on the effects of large herbivores on decomposition focused mainly on the role of microbes, we feel more attention needs to be paid to the role of the soil fauna in order to better understand ecosystem nutrient cycling.

Soil fauna also plays an important role in the decomposition of feces, with evidence of home-field advantage (HFA). Indeed deer feces decomposition in our study was more rapid on islands with deer (home) than on the island without (away), but only when including the effects of fauna (coarse-mesh bags, Fig. S3B). We infer that deer have a positive effect on macrofauna decomposing dung. Such a positive effect of large herbivores on this specialized fauna has been recently demonstrated in Japan where Iida, Soga, & Koike (2018) found a positive relationship between the populations of dung beetles and deer density. In our study, we demonstrated that, litter fauna, but not microorganisms, were selected for decomposition of a particular litter type. This is an important result as most of the literature on HFA only considered microorganisms, and this suggests a potential underestimation of fauna on HFA.

A large proportion of what we know on the effect of high quality litter deposition (dung and urine) by large herbivores on nutrient cycling comes from the study of domestic animals and/or grassland ecosystems (McNaughton, Banyikwa, & McNaughton, 1997; Frank & Groffman, 1998; Christenson, Mitchell, Groffman, & Lovett, 2010). We demonstrate that in the temperate forests we studied dung decomposed faster, and released a larger proportion of nitrogen, than observed for plant litter. Also, the addition of feces, whatever the mesh size, increased the rate of *Picea sitchensis* decomposition, increasing C loss by 31% and N loss by 47% (Fig. 4C and 4D). This may be explained by the presence of labile nutrients in dung, which enhance the development of microbial communities, increasing rates of nutrient cycling (Bardgett, Keiller, Cook, & Gilburn, 1998). However, despite these results, we found that high quality litter deposition did not affect overall decomposer ability (no higher decomposer ability on islands with deer dung/urine, Fig. 2C and 2G). This contradicts recent studies which demonstrated that feces deposition enhanced plant productivity and soil nutrient availability (Barthelemy, Stark, & Olofsson, 2015; Wang et al., 2018). The explanation for the lack of effect in the forests we studied likely rests with the patchy distribution of solitary deer, in contrast to herding species like reindeer or livestock, and thus reflects the patchy, and limited, amounts of dung deposited locally, amounts that appear not to be sufficient to influence the nutrient cycling at the ecosystem level (Pastor, Naiman, Dewey, & McInnes, 1988).

## Conclusion

Our results show that in temperate forests abundant deer can play an important role in ecosystem functioning, modifying aboveground, as well as belowground, characteristics, and reducing nutrient cycling. In the last few decades, the awareness and knowledge of the effects of overabundant dee on aboveground communities has been growing worldwide (Côté et al., 2004; Takatsuki, 2009). Our study suggests that these aboveground changes are probably at the root of major modifications in nutrient cycling in temperate forest ecosystems. In addition, it has to be emphasized that our results are likely an underestimation of effects as we did not take into account the dramatic effect deer have on the quantity of litter reaching the forest floor. For example, in Western Europe the current 10 million roe deer *Capreolus capreolus*, in addition to the increasing populations of other ungulates, represent a standing biomass estimated at 0.75 billion kg that consumes ≈20 million tons of green vegetation each year (Apollonio et al., 2010). Consequently, there is critical need to expand our results to other temperate forests to assess the overall consequences increasing deer populations have on broad scale nitrogen cycling in soils (Hobbie & Villéger, 2015) and their potential influence on global carbon storage (Tanentzap & Coomes, 2012).

## Data accessibility

Data are available online: http://doi.org/10.5281/zenodo.3474045

## Acknowledgements

We want to thank Catch Catomeris, Maria Continentino, Yonadav Anbar, Barb Roswell and Max Bullock for their support in the field. This research was financially supported by the France Canada Research Fund (FCRF), University of Rennes 1 (“Défis scientifiques émergents”), the French Embassy in Canada, the French consulate in Vancouver and the Mitacs Globalink Research Award. The Research Group on Introduced Species provided financial and logistic support. The Laskeek Bay Conservation Society provided logistic support as did many members of the Haida Gwaii communities. Version 3 of this preprint has been peer-reviewed and recommended by Peer Community In Ecology (https://doi.org/10.24072/pci.ecology.100031).

## Statement of authorship

SC conceived the ideas and designed the methodology in close interaction with MM; SC, MM, JS and J-LM collected the data; MM and SC analyzed the data; SC and MM led the writing of the manuscript. All authors contributed critically to the drafts and gave final approval for publication.

## Supporting Information

**Figure S1.**
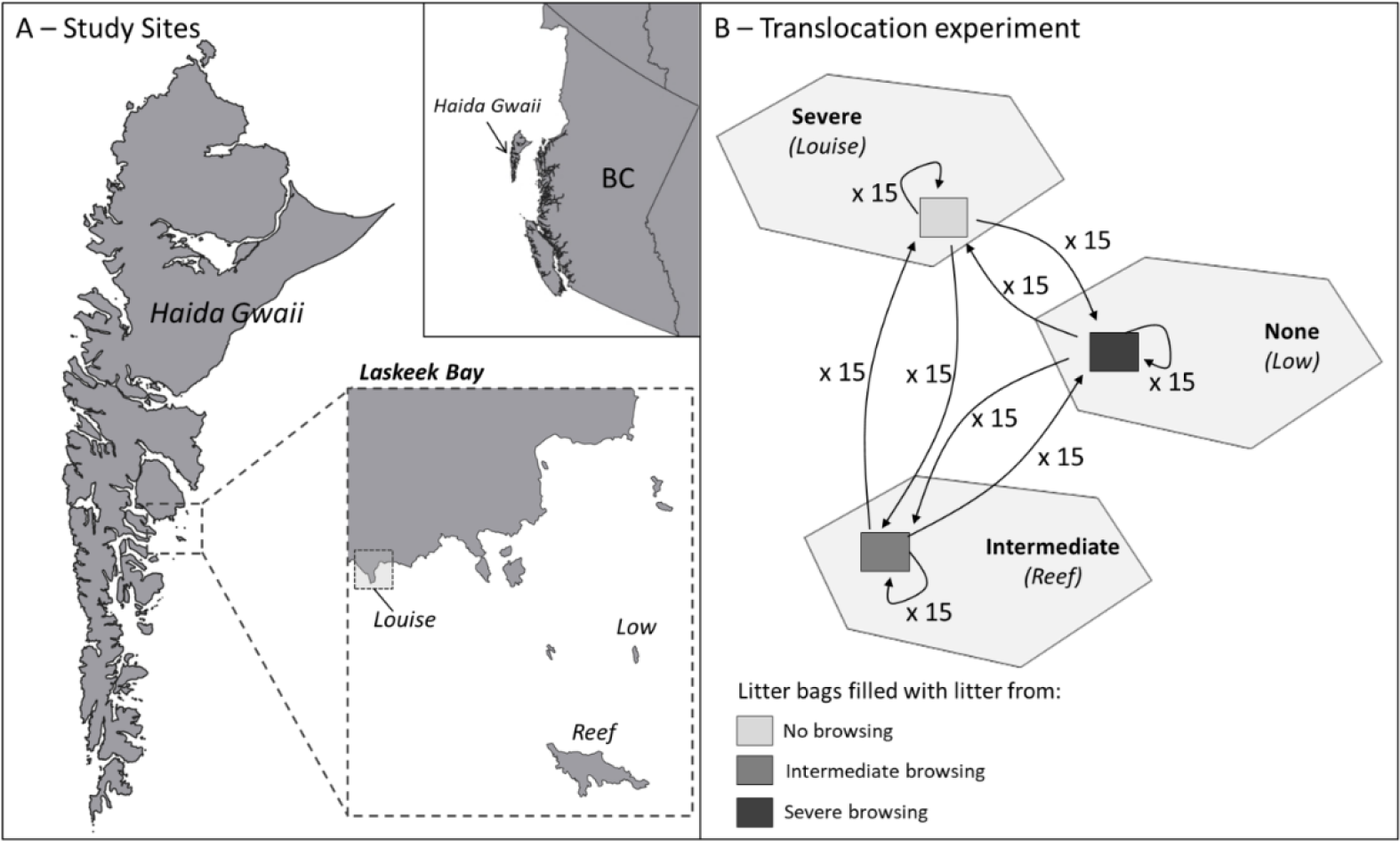
Study area and experimental design. A – Map of the study sites, B – Translocation pattern in the experiment 1. None = no deer browsing, intermediate browsing = deer present for over 70 years but exposed to significant culls between 1997 and 2010, severe browsing = deer present for over 70 years but not exposed to culls nor hunting

**Table S1.** Table synthesizing previously published results on the effect of deer on aboveground ecology of Haida Gwaii on islands covering the entire range of island sizes present in the archipelago

| Reference | Method/protocol | Main results |
| --- | --- | --- |
| **Martin et al 1995,**<br>Oikos | Songbird and vegetation sampling, 65 islands ranging from 1 to >300,000 ha | Except for the smallest most remote islands never colonized by deer, deer presence is the key factor explaining plant and animal distribution and community structure |
| **Engelstoft 1995,**<br>Master's thesis | Vegetation sampling on Graham (6,361 km<sup>2</sup>) and Moresby Islands (3,399 km<sup>2</sup>) | Deer have dramatically reduced the understory vegetation and keep the sparse understory from recovering. Deer will also have profound affects on the overstory by eliminating recruitment of Western Redcedar |
| **Martin & Baltzinger 2002,** Can. J. For. Res. | Graham (6,361 km<sup>2</sup>) and Moresby Islands (3,399 km<sup>2</sup>): in different contexts of deer hunting pressure | Regeneration of western redcedar (*Thuja plicata*) is drastically reduced in presence of deer |
| **Allombert et al. 2005,**<br>Conservation Biology | Six small islands of Laskeek Bay with different browsing histories (no deer vs deer present) | Insect abundance in the vegetation decreased eightfold and species density sixfold on islands with deer |
| **Stockton et al. 2005,**<br>Biological Conservation | Seven small islands of Laskeek Bay with different browsing histories (no deer vs deer present) | Vegetation cover exceeded 80% in the lower vegetation layers on islands without deer and was less than 10% on the islands with deer |
| **Gaston et al. 2006,**<br>Ecoscience | Ten islands of Laskeek Bay with different browsing histories (no deer vs deer present) ranging from 4.5 to 395 ha | Reversal of the normal species number-island area relationship as a result of deer browsing. Conclude that deer are a major factor structuring the island plant communities |
| **Stroh et al. 2008,** Forest Ecology & Management | Graham (6,361 km<sup>2</sup>): deer exclosure | Protected seedlings survived better, were higher, presented more leafed shoots, and had less stems than unprotected individuals |
| **Chollet et al. 2015,**<br>Biological Invasions | 57 islands ranging from 1 to 425 ha with different browsing histories | Deer are the main factor explaining the abundance of understory vegetation and understory songbirds on the islands except for the few small isolated islands never colonized by deer. |

**Table S2.**
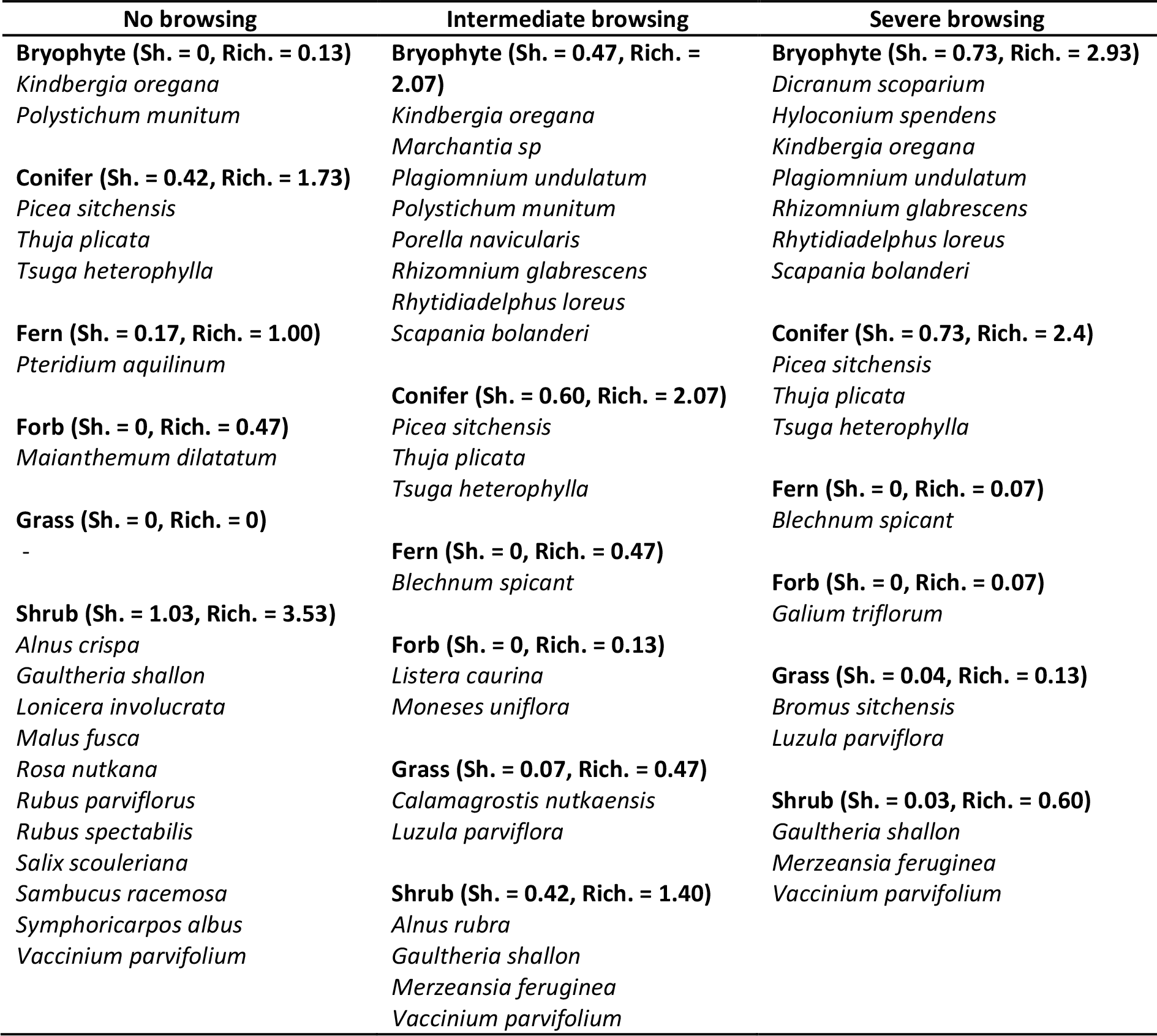
List of plant species recorded in the three browsing treatments. All species had a percent cover higher than 5 % in at least one plot. Mean shannon index (Sh.) and richness (Rich.) are given for each plant guild. Mean percent covers are given for each species.

**Table S3.** ANOVA tables of the models explaining the mass, carbon and nitrogen loss according to the litter composition and the decomposition place.

| Model | F-value | p-value |
| --- | --- | --- |
| **Mass loss in fine-mesh bags ~** |  |  |
| Composition | 117.36 | **<0.001** |
| Decomposition place | 3.02 | **0.05** |
| Composition * Decomposition place | 1.08 | 0.37 |
| **Mass loss in coarse-mesh bags ~** |  |  |
| Composition | 9.49 | **<0.001** |
| Decomposition place | 4.54 | **0.01** |
| Composition * Decomposition place | 0.80 | 0.53 |
| **C loss ~** |  |  |
| Composition | 108.78 | **<0.001** |
| Decomposition place | 2.44 | 0.09 |
| Composition* Decomposition place | 1.03 | 0.40 |
| **N loss ~** |  |  |
| Composition | 17.53 | **<0.001** |
| Decomposition place | 1.15 | 0.32 |
| Composition * Decomposition place | 0.03 | 1 |

**Table S4.** ANOVA tables of the models explaining carbon and nitrogen loss according to the decomposition place and the CWM litter C:N.

| Model | F-value | p-value | R<sup>2</sup> |
| --- | --- | --- | --- |
| C loss ~ |  |  |  |
| CWM | 11.49 | **9.4 e<sup>-4</sup>** | 0.098 |
| Decomposition place | 0.99 | 0.38 |  |
| CWM* Decomposition place | 0.09 | 0.92 |  |
| N loss ~ |  |  |  |
| CWM | 119.75 | **2e<sup>-16</sup>** | 0.505 |
| Decomposition place | 1.77 | 0.18 |  |
| CWM* Decomposition place | 0.96 | 0.38 |  |

**Figure S2.**
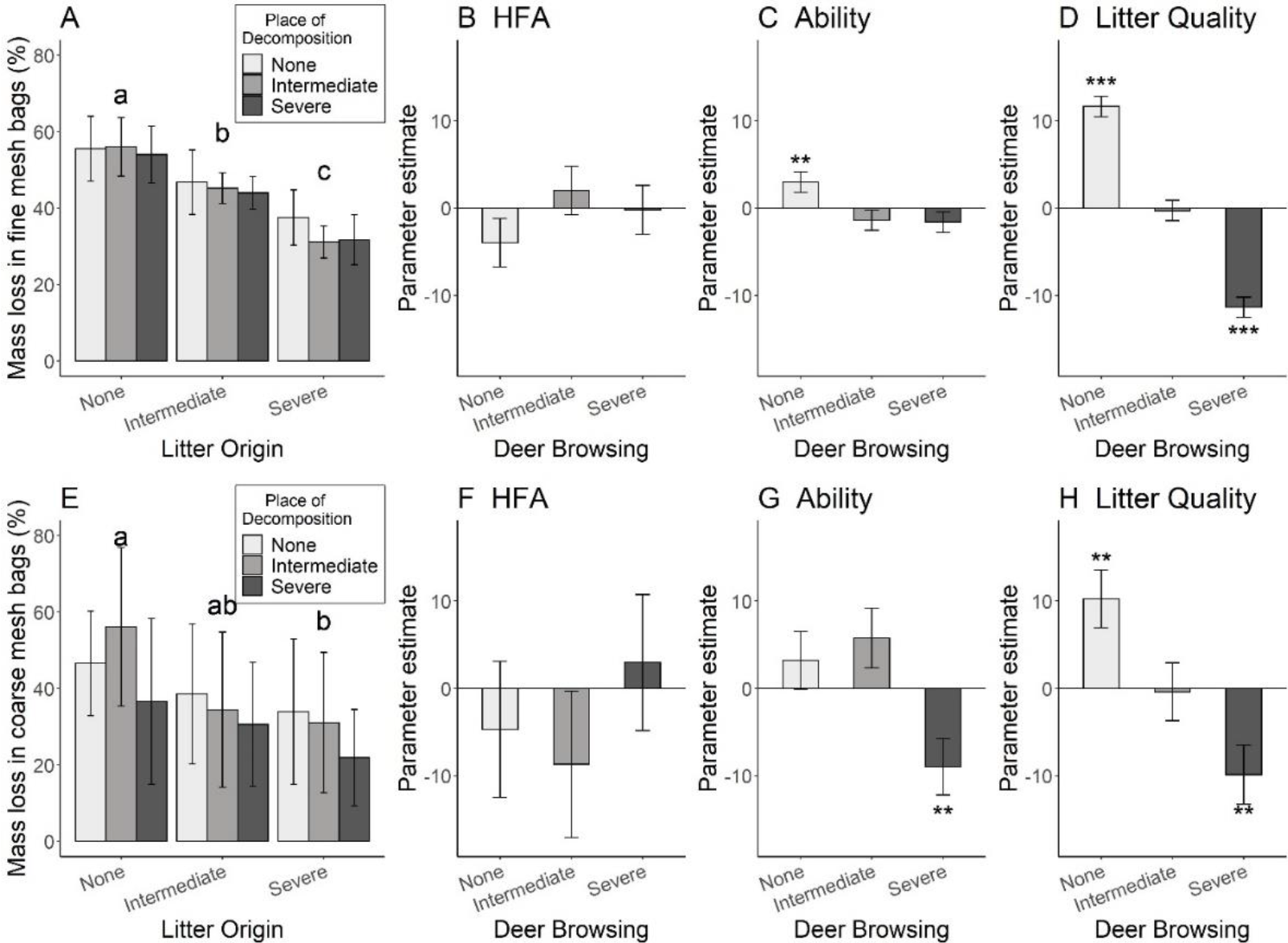
Mass loss after one year of the plant litter among herbivory treatments for fine-mesh litterbags (top) and coarse-mesh litterbags (bottom) observed in the translocation experiment. Shades of barplots represent the herbivory treatment with: light grey = no browsing (no deer), grey = intermediate (deer present for over 70 years but exposed to significant culls between 1997 and 2010) and dark grey = severe browsing intensity (deer present for over 70 years but not exposed to hunting). Asterisks indicate estimates significantly different from zero with *<0.05, ** <0.01, ***<0.001. Panel A and Panel E represent mass loss among treatments in fine and coarse-mesh litter bags respectively with bars grouped according to litter origin (X axis) and shades corresponding to the place of decomposition. Panel B to D and F to G represent the parameter estimates (± SE) calculated using the Decomposer Ability Regression Test proposed by Keiser et al. (2014)

**Figure S3.**
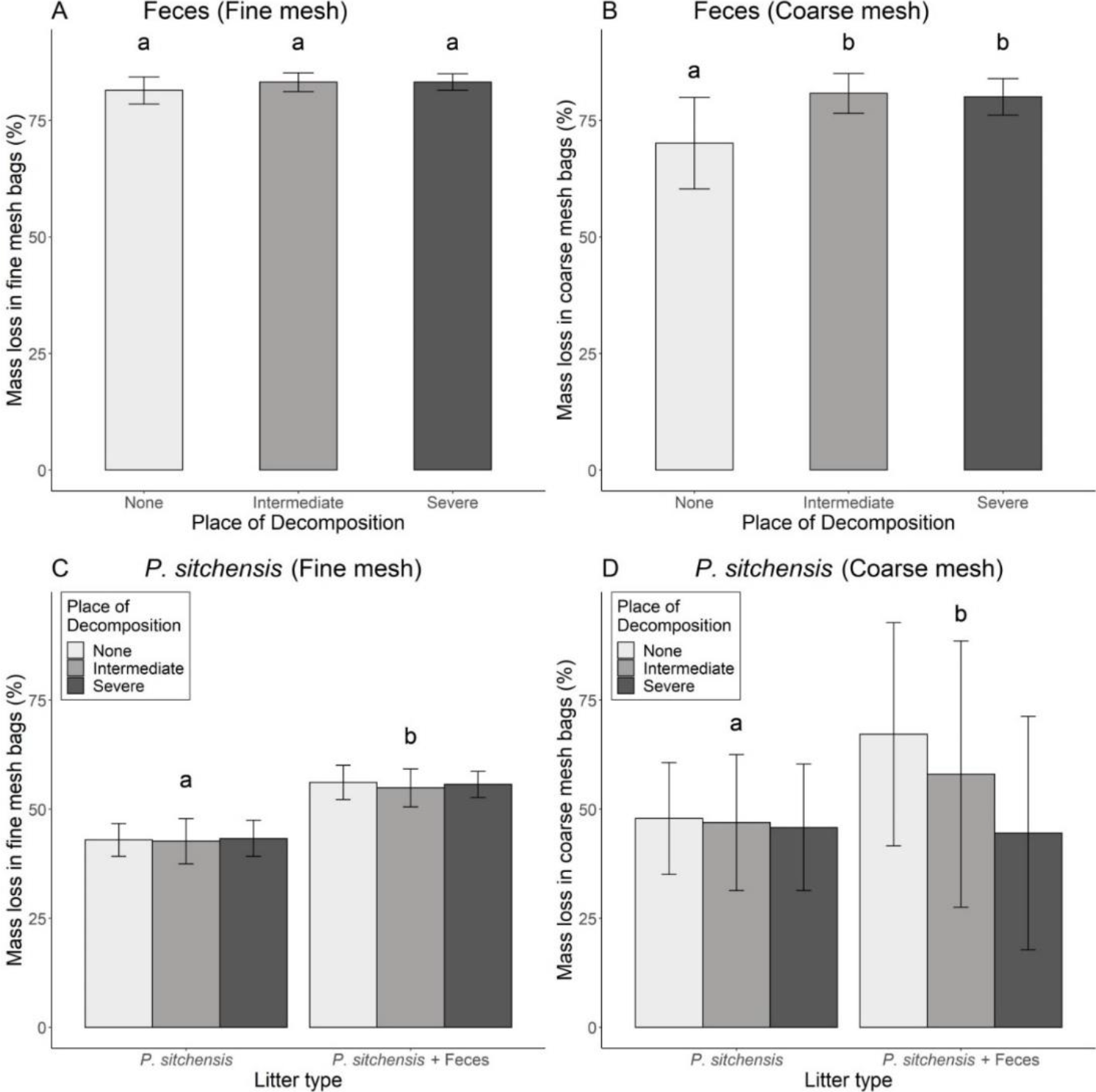
Decomposition of feces (top) and effect of feces addition on *Picea sitchensis* decomposition (bottom) for fine-mesh (left) and coarse-mesh (right) litterbags. Shades of barplots represent the browsing intensity of the place of decomposition with: light grey = no browsing (no deer), grey = intermediate (deer present for over 70 years but exposed to significant culls between 1997 and 2010) and dark grey = severe browsing (deer present for over 70 years but not exposed to hunting). Panel A – Mass loss in feces in fine-mesh litter bags; Panel B – Mass loss in feces in coarse-mesh litter bags; Panel C – Mass loss in *P. sitchensis* litter with and without the addition of feces in fine-mesh litter bags; Panel D – Mass loss in *P. sitchensis* litter with and without the addition of feces in coarse-mesh litter bags.

